# Putative rapid-acting antidepressant nitrous oxide (“laughing gas”) evokes rebound emergence of slow EEG oscillations during which TrkB signaling is induced

**DOI:** 10.1101/282954

**Authors:** Samuel Kohtala, Wiebke Theilmann, Marko Rosenholm, Paula Kiuru, Salla Uusitalo, Kaija Järventausta, Arvi Yli-Hankala, Jari Yli-Kauhaluoma, Henna-Kaisa Wigren, Tomi Rantamäki

## Abstract

Electroconvulsive therapy (ECT) remains among the most efficient antidepressants but it seldom brings immediate remedy. However, a subanesthetic dose of NMDA-R (N-methyl-D-aspartate receptor) blocker ketamine ameliorates symptoms of depression already within hours. Glutamatergic excitability and regulation of TrkB neurotrophin receptor and GSK3β (glycogen synthase kinase 3β) signaling are considered as molecular-level determinants for ketamine’s antidepressant effects. Recent clinical observations suggests that nitrous oxide (N_2_O, “laughing gas”), another NMDA-R blocking dissociative anesthestic, also produces rapid antidepressant effects but the underlying mechanisms remain essentially unstudied. In this animal study we show that N_2_O, with a clinically relevant dosing regimen, evokes an emergence of rebound slow EEG (electroencephalogram) oscillations, a phenomenon considered to predict the efficacy and onset-of-action ECT. Very similar rebound slow oscillations are induced by subanesthetic ketamine and flurothyl (a treatment analogous to ECT). These responses become best evident upon drug withdrawal, i.e. after the peak of acute pharmacological actions, when their most prominent effects on cortical excitability have subsided. Most importantly, TrkB and GSK3β signaling remain unchanged during N_2_O administration (ongoing NMDA-R blockade) but emerge gradually upon gas withdrawal along with increased slow EEG oscillations. Collectively these findings reveal that rapid-acting antidepressants produce cortical excitability that triggers “a brain state” dominated by ongoing slow oscillations, sedation and drowsiness during which TrkB and GSK3β signaling alterations are induced.

## Introduction

Major depression is a highly disabling psychiatric condition, the most significant risk factor for suicide and one of the biggest contributors to the global disease burden^1^. Many patients respond poorly to standard antidepressants, and with those who do respond, the therapeutic effects become evident with a delay of weeks. The huge unmet medical need for better antidepressants is well evidenced by the ongoing medical use of electroconvulsive therapy (ECT). An electric current leading into a short epileptiform EEG (electroencephalogram) activity is delivered during ECT under light anesthesia, but how this “seizure” leads into a remedy remains poorly understood. The therapeutic effects of ECT become evident faster than those of conventional antidepressants^2^, yet reduction of symptoms after a single ECT treatment is only seldom reported^3–5^. Notably, postictal emergence of slow EEG oscillations and burst suppression pattern have been associated with the efficacy and onset-of-action of ECT^6,7^. This encouraged clinical investigations to test whether deep burst-suppressing anesthesia can rapidly ameliorate depressive symptoms. Despite promising early observations the findings remained inconsistent^8–10^.

The remarkable ability of ketamine, a dissociative anesthetic and a drug of abuse, to ameliorate the core symptoms of depression already within hours after a single subanesthetic dose has stimulated great enthusiasm among scientists and clinicians^11,12^. Reported response rates to ketamine are impressive, yet many patients remain treatment-refractory^11^. Therefore, extensive efforts have been invested to find predictive efficacy markers and to uncover the precise pharmacological basis governing ketamine’s antidepressant effects. Although traditionally categorized as a non-competitive NMDA-R (N-methyl-D-aspartate receptor) blocker, ketamine has a rich pharmacodynamic profile and it regulates a large number of targets. Among them the AMPA-R (α-amino-3-hydroxy-5-methyl-4-isoxazolepropionic acid receptor) has received considerable recent attention. Emerging evidence suggests that ketamine causes surges in glutamate release and enhances AMPA-R function, which in turn augments synaptic plasticity through the BDNF (brain-derived neurotrophic factor) receptor TrkB^13–16^. Inhibition of GSK3β (glycogen synthase kinase 3β), another molecular event tightly connected with ketamine’s therapeutic effects^17^, contributes to enhanced AMPA-R function^18^. Most interestingly, a recent preclinical report suggests that a metabolic byproduct of ketamine, cis-6-hydroxynorketamine (HNK), directly promotes AMPA-R function and thereby governs the antidepressant effects of ketamine^19^. This hypothesis, however, conflicts with numerous earlier investigations emphasizing the critical role of NMDA-R blockade and the promising clinical observations with some other NMDA-R antagonists in depressed patients^20^. Of these agents nitrous oxide^21^ (N_2_O, “laughing gas”) is particularly interesting since (in both mice and humans) it has extremely fast kinetics and is essentially not metabolized. Notably, in the clinical study conducted by Nagele and collegues^21^ the antidepressant effects of N_2_ O was observed an hour after the gas administration, a time period when the drug has been completely eliminated from the body.

To provide better understanding on the rapid antidepressant effects we investigated how N_2_O regulates brain activity measured by the EEG and the key “molecular determinants” implicated in rapid antidepressant mechanisms in adult mice during gas exposure and withdrawal. Our findings suggests that N_2_O, similarly to that seen with subanesthetic ketamine and volatile convulsant flurothyl (analogous with ECT), produces transient cortical excitation resulting into rebound emergence of slow EEG oscillations. Most interestingly, TrkB and GSK3β signaling alterations remain unchanged during N_2_O exposure but are evoked gradually upon drug withdrawal along with slow oscillations.

## Materials and Methods

### Animals

Adult C57BL/6JRccHsd mice (Harlan Laboratories, Venray, Netherland) were used. Animals were maintained in the animal facility of University of Helsinki, Finland, under standard conditions (21 °C, 12-hour light-dark cycle) with free access to food and water. The experiments were carried out according to the guidelines of the Society for Neuroscience and were approved by the County Administrative Board of Southern Finland (License: ESA-VI/10527/04.10.07/2014).

### Pharmacological treatments

Medical grade N_2_O (Livopan 50% N_2_O/O_2_ mix, Linde Healthcare; Niontix 100% N_2_O, Linde Healthcare). Medical grade oxygen (Conoxia 100% O_2_, Linde Healthcare) was mixed with 100% N_2_O to achieve >50 (75%) N_2_O concentrations. After habituation to the experimental conditions, the gas was administered into airtight Plexiglass chambers (for biochemical analyses (width × length × height): 14 cm × 25 cm × 9 cm; for biochemical & EEG analyses: 11.5 cm × 11.5 cm × 6.5 cm) with a flow rate of 4-8 l/min. O_2_ or room air was used as control gas (e.g. for the sham animals).

To induce myoclonic seizures, 10% flurothyl liquid (in 90% ethanol; Sigma-Aldrich) were administered into the cotton pad placed inside the lid of an airtight Plexi-glass chamber (13 cm × 13 cm × 13 cm) at the flow rate of 100-200 µl/min until the mice exhibited seizures. The lid was removed to terminate the seizure. Animals were euthanized at indicated times post-seizure. Ethanol solution was given for the sham animals.

Ketamine-HCl (10-200 mg/kg; Ketaminol^®^, Intervet International B.V.), 6,6-d_2_-ketamine-HCl (see below; 100 mg/kg) and cis-6-hydroxynorketamine-HCl (10-20 mg/kg; Tocris) were diluted in isotonic saline solution and injected intraperitoneally with an injection volume of 10 ml/kg.

### Synthesis of 6,6-d2-ketamine-HCl

### General information

^1^H NMR and ^13^C NMR spectra in CDCl_3_ or CD_3_OD at ambient temperature were recorded on a Bruker Avance 400 MHz NMR with the smart probe. Chemical shifts (δ) are given in parts per million (ppm) relative to the NMR reference solvent signals (CDCl_3_: 7.26 ppm; CD_3_OD: 3.31 ppm for ^1^H NMR and CDCl_3_: 77.16 ppm; CD_3_ OD: 49.0 ppm for ^13^C NMR). Multiplicities are indicated by s (singlet), br s (broad singlet), d (doublet), dd (doublet of doublets), ddd (doublet of doublet of doublets), t (triplet), m (multiplet). The coupling constants J are quoted in Hertz (Hz). HRMS spectra were recorded using Waters Acquity UPLC^®^-system (with Acquity UPLC^®^ BEH C18 column, 1.7 µm, 50 × 2.1 mm, Waters) with Waters Synapt G2 HDMS with the ESI (+), high resolution mode. The mobile phase consisted of H_2_O (A) and acetonitrile (B) both containing 0.1% HCOOH. Microwave syntheses were performed in sealed tubes using Biotage Initiator+ instrument equipped with an external IR sensor.

Racemic ketamine-HCl **(1)** (0.60 g, 2.2 mmol (YA Apteekki, Helsinki, Finland) was dissolved in dry THF (6 mL) and D_2_O (2.25 mL). To this solution a 40% solution of NaOD in D_2_O (2.25 mL) was added. The sealed tube was microwave-irradiated at 120 °C for 2 h. The resulting mixture was poured to 1 M aqueous solution of HCl (20 mL). The white precipitate was filtered and washed with water and dried to yield 6,6-d_2_ -ketamine **(2)** (394 mg, 75%). ^1^H NMR and MS spectra showed that the isotopic purity was not >90%, so the above procedure was repeated (379 mg) using a 40% solution of NaOD in D_2_O (1 mL). 6,6-d_2_-Ketamine **(2)** was obtained as a white solid (342 mg, 90%). ^1^H NMR (400 MHz, CDCl_3_) δ 7.55 (dd, J = 7.9, 1.7 Hz, 1H), 7.38 (dd, J = 7.8, 1.5 Hz, 1H), 7.32 (ddd, J = 7.8, 7.3, 1.5 Hz, 1H), 7.27–7.21 (m, 1H), 2.84–2.73 (m, 1H), 2.15 (bs, 1H), 2.10 (s, 3H), 2.03-1.95 (m, 1H), 1.90-1.80 (m, 1H), 1.74 (m, 3H). ^13^C NMR (101 MHz, CDCl_3_) δ 209.5, 137.9, 134.0, 131.4, 129.6, 128.8, 126.8, 70.4, 39.1 (m), 38.8, 29.3, 28.2, 22.0. 6,6-d_2_-Ketamine (0.30 g was dissolved in dry 1,4-dioxane (5 mL) and a 4 M solution of HCl in 1,4-dioxane (1 mL) was added. The formed white precipitate was filtered and dried to yielding 6,6-d_2_-ketamine-HCl **(3)** (0.26 g, 75%). 6,6-d_2_-ketamine-HCl was dissolved to water, filtered and dried prior the use.

^1^H NMR (400 MHz, CD_3_OD) δ 7.94–7.90 (m, 1H), 7.68–7.58 (m, 3H), 3.42–3.34 (m, 1H), 2.39 (s, 3H), 2.17– 2.10 (m, 1H), 1.96–1.84 (m, 2H), 1.83–1.68 (m, 2H), ^13^C NMR (101 MHz, CD_3_OD) δ 208.4, 135.9, 133.9, 133.4, 133.2, 129.7, 129.2, 73.6, 40.3 (m), 37.4, 30.9, 27.9, 22.8. ^1^H NMR and ^13^C NMR were in agreement with reported data^19^. HRMS (ESI+): Calcd for C_13_H_15_D_2_ClNO^+^ 240.1124; found 240.1130.

### Western blotting and quantitative RT-PCR

Animals were sacrificed at indicated times after the treatments by rapid cervical dislocation followed by decapitation. No anesthesia was used due to its potential confounding effects on the analyses^22^. Bilateral medial prefrontal cortex (including prelimbic and infralimbic cortices) was rapidly dissected on a cooled dish and stored at -80 °C^22,23^.

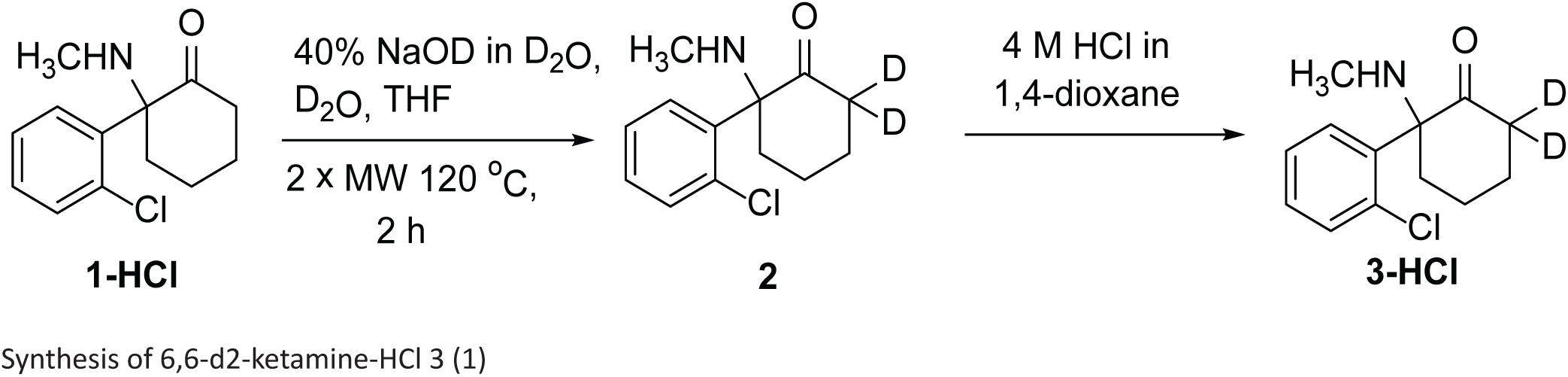

For western blotting the brain samples were homogenized in lysis buffer (137 mM NaCl, 20 mM Tris, 1% NP-40, 10% glycerol, 48 mM NaF, H_2_O, Complete inhibitor mix (Roche), PhosStop (Roche))^23^. After ∼15 min incubation on ice, samples were centrifuged (16000g, 15 min, +4 °C) and the resulting supernatant collected for further analysis. Sample protein concentrations were measured using Bio-Rad DC protein assay (Bio-Rad Laboratories, Hercules, CA). Proteins (40-50 μg) were separated with SDS-PAGE under reducing and denaturing conditions and blotted to a PVDF membrane as described^22,23^. Membranes were incubated with the following primary antibodies (see Ref.^22^): anti-p-TrkB (#4168; 1:1000; Cell signaling technology (CST)), anti-TrkB (1:1000; #4603, CST), anti-Trk (sc-11; 1:1000; Santa Cruz Biotechnology (SCB);), anti-p-CREB (#9191S; 1:1000; CST), anti-p-p70S6K (#9204S; 1:1000; CST), anti-p-GSK3βS9 (#9336; 1:1000; CST), anti-p-p44/42-MAPK^Thr202/Y204^ (#9106, 1:1000, CST), anti-GSK3β (#9315, 1:1000, CST), anti-p70S6K (#2708, 1:1000, CST) anti-p44/42-MAPK (#9102, 1:1000, CST) and anti-GAPDH (#2118, 1:10 000, CST). Further, the membranes were washed with TBS/0.1% Tween (TBST) and incubated with horseradish peroxidase conjugated secondary antibodies (1:10000 in non-fat dry milk, 1 h at room temperature; Bio-Rad). After subsequent washes, secondary antibodies were visualized using enhanced chemiluminescence (ECL Plus, ThermoScientific, Vantaa, Finland) for detection by Biorad ChemiDoc MP camera (Bio-Rad Laboratories, Helsinki, Finland).

For qPCR, total RNA of the sample was extracted using Trizol (Thermo Scientific) according to the manufacturer’s instructions and treated with DNAse I mix. mRNA was reverse transcribed using oligo (dT) primer and SuperScript III Reverse Transcriptase mix (Thermo Scientific). The amount of cDNA was quantified using real-time PCR. The primers used to amplify specific cDNA regions of the transcripts are shown in **Supplementary Table 1**. DNA amplification reactions were run in triplicate in the presence of Maxima SYBRGreen qPCR mix (Thermo Scientific). Second derivate values from each sample were obtained using the LightCycler 480 software (Roche). Relative quantification of template was performed as described previously using standard curve method, with cDNA data being normalized to the control *Gapdh* and *ß-actin* level.

### EEG recordings and data analysis

For the implantation of electrodes, mice were anesthetized with isoflurane (3% induction, 1.5-2% maintenance). Lidocaine was used as local anesthetic and buprenorphine (0.1 mg/kg, s.c.) for postoperative care. Two epidural screw EEG (electroencephalogram) electrodes were placed above the fronto-parietal cortex. A further screw served as mounting support. Two silver wire electrodes were implanted in the nuchal muscles to monitor the EMG (electromyogram). After the surgery, mice were single-housed in Plexiglas boxes. After a recovery period of 5-7 days, animals were connected to flexible counterbalanced cables for EEG/EMG recording and habituated to recording cables for three days.

Baseline EEG (∼10 min) recordings of awake animals were conducted prior the treatments. All injection treatments were conducted in the animal’s home cages during light period. N_2_O treatment was delivered in home-made anesthesia boxes for indicated time periods with a flow rate of 8 l/min.

The EEG and EMG signals were amplified (gain 5 or 10 K) and filtered (high pass: 0.3 Hz; low pass 100 Hz; notch filter) with a 16-channel AC amplifier (A-M System, model 3500), sampled at 254 Hz or 70 Hz with 1401 unit (CED), and recorded using Spike2 (version 8.07, Cambridge Electronic Devices). The processing of the EEG data was obtained using Spike2 (version 8.07, Cambridge Electronic Devices). EEG power spectra were calculated within the 1-50 Hz frequency range by fast Fourier transform (FFT = 256, Hanning window, 1.0 Hz resolution). Oscillation power in each bandwidth (delta=1–4 Hz; theta=4–7 Hz; alpha=7–12 Hz; beta=12–25 Hz; gamma low=25–40 Hz; gamma high=60-100 Hz) was computed in 30-300-sec epochs from spectrograms (FFT size: 1024 points) for each animal. Representative sonograms were computed using a Hanning window with a block size of 512.

### Statistical analyses

Depending on whether data were normally distributed or not, either parametric or nonparametric tests were used for statistical evaluation. In case of more than two groups, analysis of variance (ANOVA) with post hoc test was used. All statistical analyses were performed with the Prism 7 software from GraphPad (La Jolla, CA, USA). All tests were two-sided; a P≤0.05 was considered significant. Details of statistical tests and n numbers for each experiment are shown in **Supplementary Table 2**.

## Results

To provide further insights into the putative antidepressant effects of N_2_O, and to identify potential shared mechanistic principles among diverse treatments carrying rapid antidepressant potential, we performed pharmaco-EEG recordings (at bandwidth of ∼1-100 Hz) in freely moving mice subjected to flurothyl, subanesthetic ketamine and subanesthetic N_2_O treatments. We first adopted the N_2_O treatment protocol used in the clinical study by Nagele et al^21^ (50% N_2_O for 60 min) while a single intraperitoneal dose of 10 mg/kg was selected for ketamine based on previous animal experiments^15,24^. Flurothyl was evaporated into the cage until the mice exhibited a generalized seizure; which terminated within seconds upon drug withdrawal.

A robust increase in slow EEG oscillations, particularly within the delta range (1-4 Hz), emerged gradually and peaked within 10 min after the flurothyl-induced seizure **(Figure 1A)**. Alpha oscillations (7-12 Hz), beta oscillations (12-30 Hz) and high frequency gamma oscillations (>25 Hz) were reduced during the post-ictal (i.e. after seizure) period. At the behavioural level the mice appeared motionless and sedated, a state also correlated with reduced electromyogram (EMG) activity **(Figure 1A)**. As reported earlier^19,25^, subanesthetic ketamine initially increased gamma oscillations **(Figure 1B)**, a neurophysiological sign of increased cortical excitability, that lasted around 30-50 min. At this time period, i.e. after the peak of ketamine’s pharmacological effects (serum t_½_ (mouse): ∼15 min, see Ref. ^26^), slow-wave delta oscillations gradually increased above baseline and saline treated controls **(Figure 1B; see also Supplementary Figure 1)**. Apart from the dampening of low gamma oscillations, no clear EEG alterations were observed during N_2_O exposure **(Figure 1C)**. Upon gas withdrawal, however, slow EEG oscillations increased above baseline values. The peak of slow-wave delta emerged at around 40 minutes post-N_2_O and reduced thereafter towards baseline. Taking into account the pharmacokinetics of the drugs these data indicate that slow EEG oscillations are triggered in the cortex as a rebound response to preceding exposure to flurothyl, subanesthetic ketamine and N_2_O. Such a phenomenon has been previously observed with ketamine, and another NMDA-R blocker MK-801, in rodents and has been suggested to occur as a homeostatic response to the transient cortical excitation induced by the drug^27,28^. Post-ictal emergence of slow EEG oscillation is observed in patients after the delivery of flurothyl or ECT^29^ and this phenomenon has been considered to predict the efficacy and onset-of-action of convulsive therapies^6,7^.

**Figure 1.**
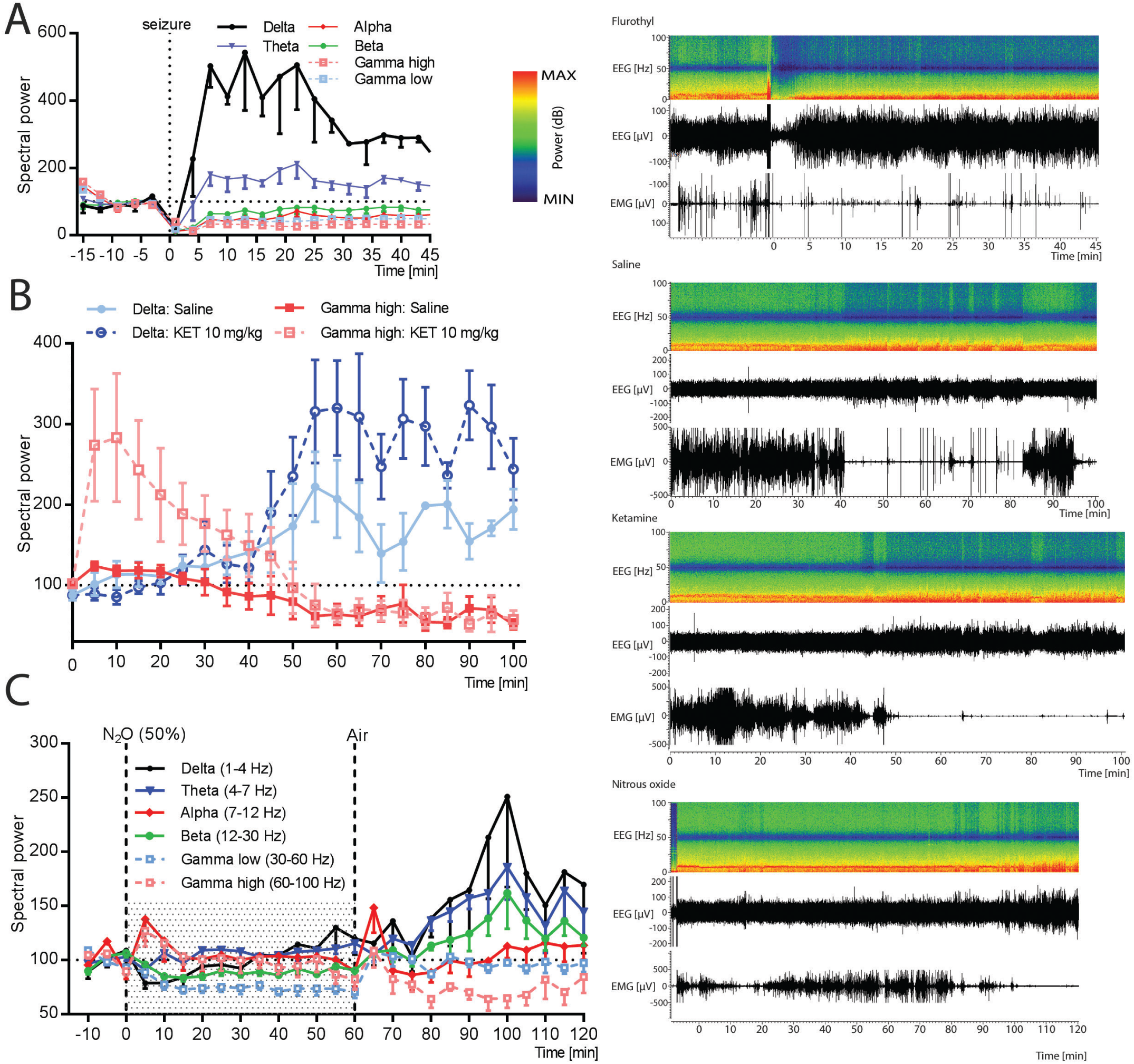
Rapid-acting antidepressants evoke homeostatic emergence of slow EEG oscillations upon drug withdrawal. **(A)** Flurothyl (FLUR) induced seizures evoke rebound emergence of slow-wave delta (1-4 Hz) and theta (4-7 Hz) oscillations. Representative time frequency EEG spectrogram and normalized power of major EEG oscillations before, during and after drug administration. **(B)** Subanesthetic ketamine (KET; 10 mg/kg, i.p.) evokes rebound delta oscillations gradually after the acute effects of the drug on high gamma oscillations have dissipated. **(C)** Representative time frequency EEG spectrogram and normalized power of major EEG oscillations before, during and after the exposure to 50% nitrous oxide (N_2_O) for 60 minutes. Data are means ± S.E.M. (for n numbers see **Supplementary Table 2**).

N_2_O and most other NMDA-R antagonists, including ketamine, are so called dissociative anesthetics and their use may promote psychoactive effects and perceptual distortions^30^. Unlike GABAergic anesthetics, cerebral metabolic rate is increased during exposure to N_2_O and ketamine^31,32^. Therefore, we next investigated the effects of N_2_O on markers implicated in cortical excitability. We focused our studies to the medial prefrontal cortex (mPFC), a brain region associated in the pathophysiology of depression and antidepressant actions. The mRNA expression levels of well known activity-dependent immediate early genes (IEGs) (*c-fos, arc, bdnf, zif-268*) were significantly increased an hour after withdrawal from 50% N_2_O (**Figure 2A)**. *Homer-1A, egr-2, mkp-1* and *synapsin* mRNAs were also increased **(Figure 2A)**. Notably, IEGs and phosphorylation of mitogen-activated protein kinase (MAP-K^T202/Y204^) were up-regulated in samples collected without a washout period **(Figure 2B)**, indicating that cortical activity is facilitated under the acute influence of N_2_O. These acute molecular effects of N_2_O resemble those produced by electroconvulsive shock (a rodent model of ECT)^33,34^ and sleep deprivation^35^, which also alleviates symptoms of depression rapidly in a subset of patients^36^.

**Figure 2.**
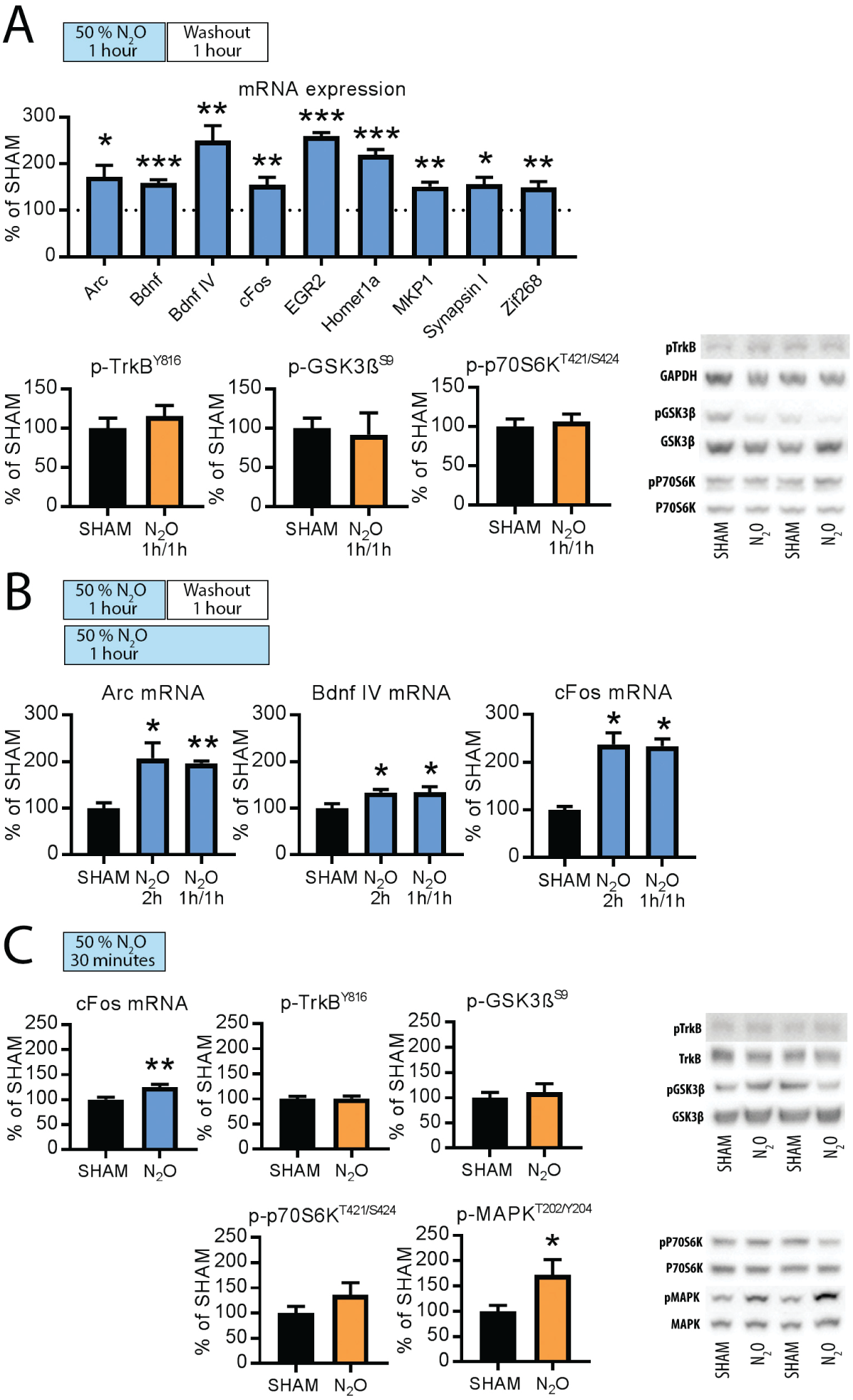
Nitrous oxide (N_2_O) increases excitation markers in the adult mouse medial prefrontal cortex. **(A)** *c-fos, arc, bdnf, egr2, homer-1a, mkp-1, zif-268* and *synapsin* mRNAs levels up-regulated 1-hour after N_2_O (50%) exposure while the levels of p-TrkB^Y816^, p-GSK3β^S9^ and p-p70S6k^T421/424^ remain unaltered. **(B)** *c-fos, arc, bdnf mRNAs* levels up-regulated during 2-hour N_2_O (50%) exposure. **(C)** *c-fos* mRNA and p-MAPK^T202/Y204^ levels up-regulated during 30-min N_2_O (50%) exposure while the levels of p-TrkB^Y816^, p-GSK3β^S9^ and p-p70S6k^T421/424^ remain unaltered. Data are means ± S.E.M. *<0.05, **<0.01, ***<0.005 (for statistical analyses and n numbers see **Supplementary Table 2**).

Among molecular level mechanisms, activation of TrkB receptor and inhibition of GSK3β kinase have been causally connected with antidepressant effects in rodents^14–17,24^. Upon activation TrkB receptors undergo tyrosine phosphorylation within intracellular domains^37^ and phosphorylation the phospholipase-Cγ1 binding site (tyrosine-816) is readily regulated by antidepressants^23^. However, phosphorylation of TrkB^Y816^ (and its down-stream kinase p70S6k^T421/S424^, Ref. 5) and phosphorylation of GSK3β at the inhibitory serine-9 residue^38^ (GSK3β^S9^) remained unaltered in samples collected an hour after 50% N_2_O treatment **(Figure 2A)**. When samples were collected and analyzed from mice euthanized during N_2_O administration similar effects were seen **(Figure 2C)**, indicating that NMDA-R blockade insufficient enough to produce anesthesia is not coupled with TrkB and and GSK3β signaling alterations. In line with this, albeit unexpectedly^16,24^, TrkB^Y816^, p70S6k^T421/S424^ and GSK3β^S9^ phosphorylation remained unaltered also 30 min after a subanesthetic dose of ketamine **(Figure 3A)**.

**Figure 3.**
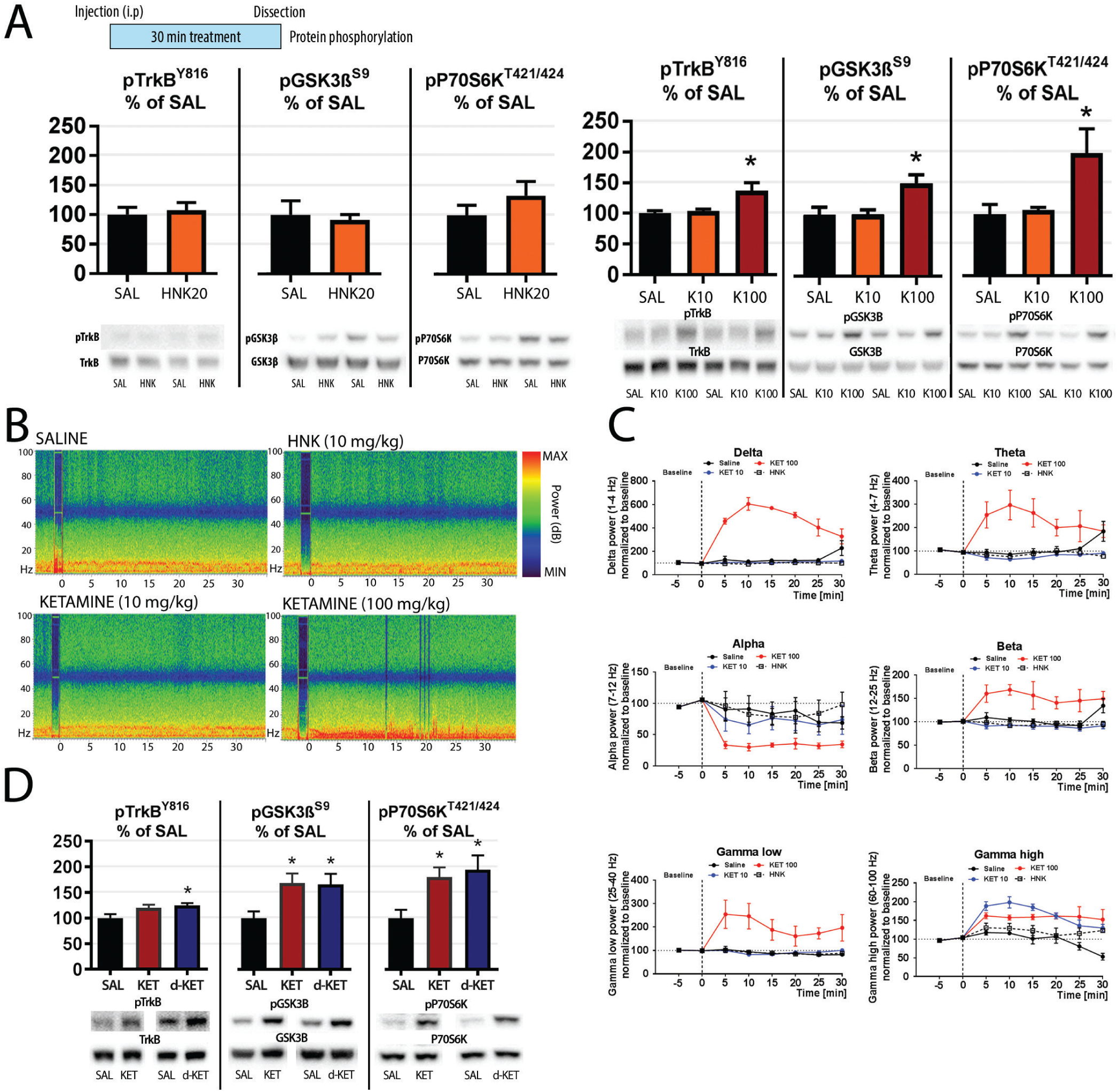
Acute effects of ketamine and cis-6-hydroxynorketamine on TrkB and GSK3β signaling and EEG. **(A)** Phosphorylation of TrkB^Y816^, GSK3β^S9^ and p70S6k^T421/424^ in the adult mouse medial prefrontal cortex 30 min after an i.p. injection of saline (SAL), cis-6-hydroxynorketamine (HNK, 20 mg/kg) or ketamine (KET, 10 mg/kg, 100 mg/kg). **(B)** Representative time frequency EEG spectrograms immediately before and during HNK and KET treatment. **(C)** Normalized power of major EEG oscillations during HNK and KET treatment (data analyzed in 5 min bins). Dashed vertical line indicates injection point (0 min). **(D)** Effects of KET (100 mg/kg, i.p.; 30 min) and 6,6-dideuteroketamine (d-KET, 100 mg/kg, i.p.; 30 min) on p-TrkB^Y816^, p-GSK3β^S9^ and p-p70S6k^T421/424^. Data are means ± S.E.M. *<0.05, **<0.01, ***<0.005 (for statistical analyses and n numbers see **Supplementary Table 2**).

A recent preclinical report suggests that HNK, a metabolic byproduct of ketamine, directly facilitates AMPA-R function and thereby governs antidepressant effects^19^. Compared to ketamine, HNK acts as a weak NMDA-R antagonist^39^ and is devoid of psychotomimetic and anesthetic properties^19^. To investigate whether drug-induced facilitation of AMPA-R function^19^ regulates TrkB and GSK3β signaling, we subjected mice to acute HNK treatments. The phosphorylation of TrkB^Y816^, p70S6k^T421/ S424^ and GSK3β^S9^ remained however again unaltered **(Figure 3A)**. Intriguingly however, the ability of ketamine to acutely regulate all these phosphorylation changes increased dose-dependently and by and large the most significant changes were seen with sedative-anesthetic doses **(Figure 3A; Supplementary Figure 2)**. At the level of EEG, these doses produced most prominent increase in slow oscillations, although all major EEG oscillations were affected **(Figure 3B-C)**. Most importantly, ketamine deuterated at the C6 position (6,6-d_2_-Ketamine), a modification that reduces its metabolism into HNK^19^, recapitulated the phosphorylation effects of an equivalent dose of ketamine on TrkB and GSK3β **(Figure 3D)**.

The proposed link between postictal emergence of slow EEG oscillations and the therapeutic effects of ECT^6,7^ and the intriguing association between anesthetic states and TrkB and GSK3β phosphorylation^22^ **(Figure 3)** prompted us next to collect brain tissues for western blot analyses 10 minutes after flurothyl-induced seizure when slow EEG oscillations are robustly elevated. Indeed, phosphorylation levels of TrkB^Y816^, p70S6k^T421/S424^ and GSK3β^S9^ were significantly increased in these samples **(Figure 4A)** indicating that TrkB and GSK3β signaling responses become effective specifically during evoked slow oscillations also associated with sedation and drowsiness. To test this hypothesis with N_2_O we first investigated whether rebound slow EEG oscillations can be more readily induced by higher N_2_O concentrations. Indeed, slow oscillations elevated within minutes after a short exposure to 75% of N_2_O **(Figure 4B; Supplementary Figure 3)**. Beta and low gamma oscillations were reduced during this treatment but these alterations normalized upon gas withdrawal **(Supplementary Figure 3)**. These data encouraged us to collect brain samples for western blot analyses during these withdrawal periods (5 or 15 min) after exposing the animals to varying N_2_O concentrations (50-75%) for 20 minutes. These data, shown in **Figure 4C-D**, reveal that N_2_O can indeed induce TrkB^Y816^ and GSK3β^S9^ phosphorylation but only upon gas withdrawal when slow EEG oscillations become facilitated.

**Figure 4.**
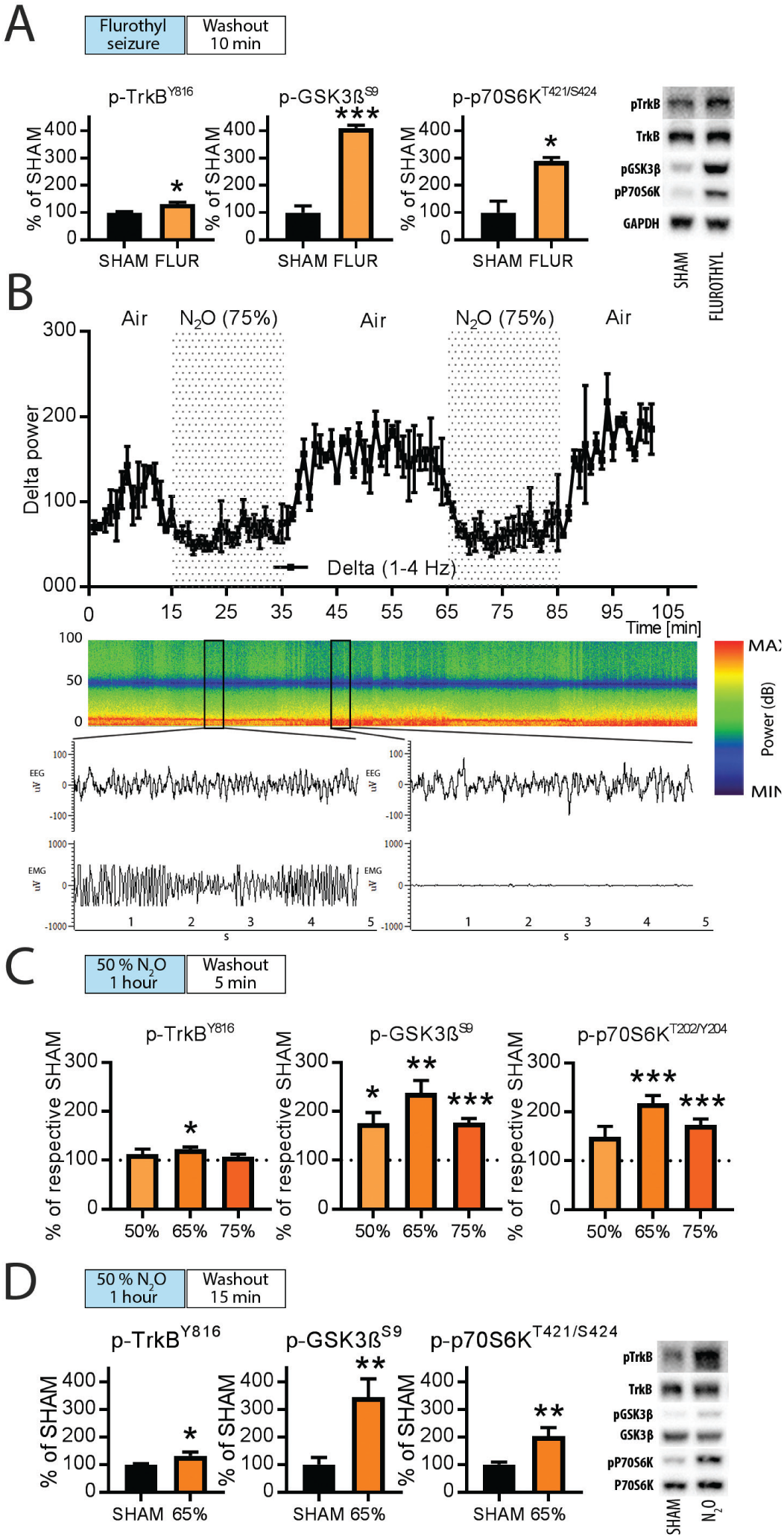
TrkB and GSK3β signaling become regulated during evoked slow EEG oscillations induced by flurothyl or nitrous oxide (N_2_ O). **(A)** Levels of p-TrkB^Y816^, p-GSK3β^S9^ and p70S6k^T421/424^ 10 min after flurothyl-induced seizure (post-ictal state). **(B)** Rebound delta oscillations after discontinuation of 75% N_2_O treatment. (C) Levels of p-TrkB^Y816^, p-GSK3β^S9^ and p70S6k^T421/424^ at 5-minute post-N_2_O exposure (50-75%). **(D)** Levels of p-TrkB^Y816^, p-GSK3β^S9^ and p70S6k^T421/424^ at 15-minute post-N_2_O exposure (65%). Data are means ± S.E.M. *<0.05, **<0.01, ***<0.005 (for statistical analyses and n numbers see **Supplementary Table 2**).

## Discussion

Major depression gives rise to significant physical and mental disability. It often involves an acute risk to commit suicide, but commonly prescribed antidepressants bring relief very slowly, if at all. While most pharmacoresistant patients respond to ECT, rapid amelioration of symptoms after a single ECT treatment is very rare. Notably, rather than mere seizure manifestation, post-ictal emergence of slow EEG oscillations have been associated with the onset-of-action of the antidepressant effects of ECT^6,7^. Unlike any other treatment, a subanesthetic dose of ketamine has been reproducibly shown to elicit rapid antidepressant effects in multiple patient trials^11,12^ and this treatment is already in off-label use in several countries. Despite recent progress the precise neurobiological basis governing rapid antidepressant effects remain obscure and debated^13,20,22^. To get further insights into this important area of study we have here investigated how N_2_O, another NMDA-R blocking dissociative anesthetis and a putative rapid-acting antidepressant^21^, regulates EEG and the “molecular determinants” implicated in rapid antidepressant mechanisms. To increase the significance of our findings we investigated subanesthetic ketamine and flurothyl in parallel.

We reveal that flurothyl, subanesthetic ketamine and subanesthetic N_2_O all produce a characteristic emergence of slow EEG oscillations. These oscillations become evident upon drug withdrawal, i.e. after the peak of acute pharmacological action, when the most prominent effects on cortical excitability have subsided. Most remarkably, key molecular-level signaling alterations, namely activation of TrkB and inhibition of GSK3β, become specifically altered during this period of slow oscillations. Our findings show that the postictal-like of state, typically associated with convulsive therapies and characterized by slow EEG oscillations, drowsiness, confusion and nausea, is also present albeit to a milder degree after excitatory treatments such as ketamine and N_2_O. These findings provide a reason to speculate that the mechanisms of rapid antidepressant treatments might be related to the combination of both excitation induced changes gene expression and the subsequent homeostatic activation of key neurotrophic signaling pathways during postictal slow EEG oscillations **(Figure 5)**.

**Figure 5.**
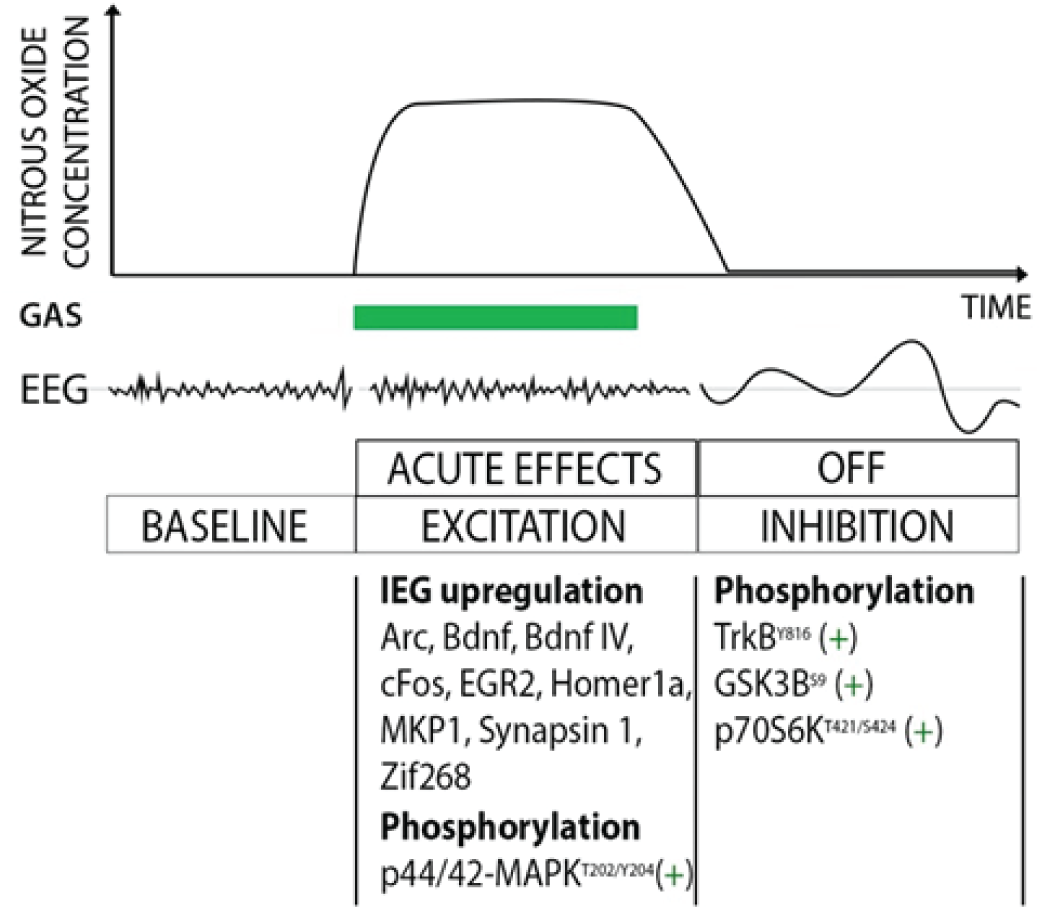
Sequential regulation of slow EEG oscillations and molecular signatures implicated in rapid antidepressant responses – nitrous oxide (N_2_O) as an example. N_2_O induces transient cortical excitability that evokes a homeostatic emergence of slow wave EEG oscillations, during which TrkB and GSK3β signaling alterations are induced. These responses become evident upon drug withdrawal, i.e. after the acute pharmacological actions (e.g. NMDA-R blockade) of N_2_O. Several biological markers implicated in cortical excitability (immediate-early genes and p42/44-MAPK) are increased during N_2_O administration.

To-date, the majority of pharmacological studies have focused to reveal specific receptor-level mediators underlying rapid antidepressant effects and less attention has been put on the crosstalk of translationally relevant network-level phenomena and molecular alterations. This report urges shifting the attention more towards the interplay of excitation and subsequent postictal state, alterations triggered within the brain as a consequence of drug challenge. That said, interconnecting the present observations with the recently identified effects of ketamine on intrinsic homeostatic plasticity processes^27,40–42^, evident as dynamic and circadian fluctuations in slow oscillations, is instrumental to provide more precise understanding of rapid antidepressant actions. The unique pharmacokinetic and pharmacological properties of N_2_O and related “fast-acting” medicines may become critical tools for these future efforts guiding the development of novel interventions against major depression.

## Acknowledgements

This study has been supported by the Academy of Finland (T.R., grants 276333, 284569, 305195, 312664) and the Finnish Funding Agency for Innovation (T.R.), the Doctoral Programme Brain & Mind (S.K.). We thank Dr. Kai Kaila, Dr. Tarja Porkka-Heiskanen, Dr. Iiris Hovatta and Dr. Raimo Tuominen for their comments on this work. Dr. Giuseppe Cortese is thanked for language editing. Tommi Makkonen, Sissi Pastell, Virpi Perko and Maria Partanen are thanked for technical assistance. Kari Tamminen (Sarlin Oy Ab) and Sari Pöysti (Oy AGA Ab) are thanked for providing equipment and advice for gas administrations.

