## Supplementary Materials for "Putative rapid-acting antidepressant nitrous oxide (“laughing gas”) evokes rebound emergence of slow EEG oscillations during which TrkB signaling is induced"

**List of Supplementary Materials**

- Supplementary Figures 1–3
- Supplementary Tables 1-2
- References

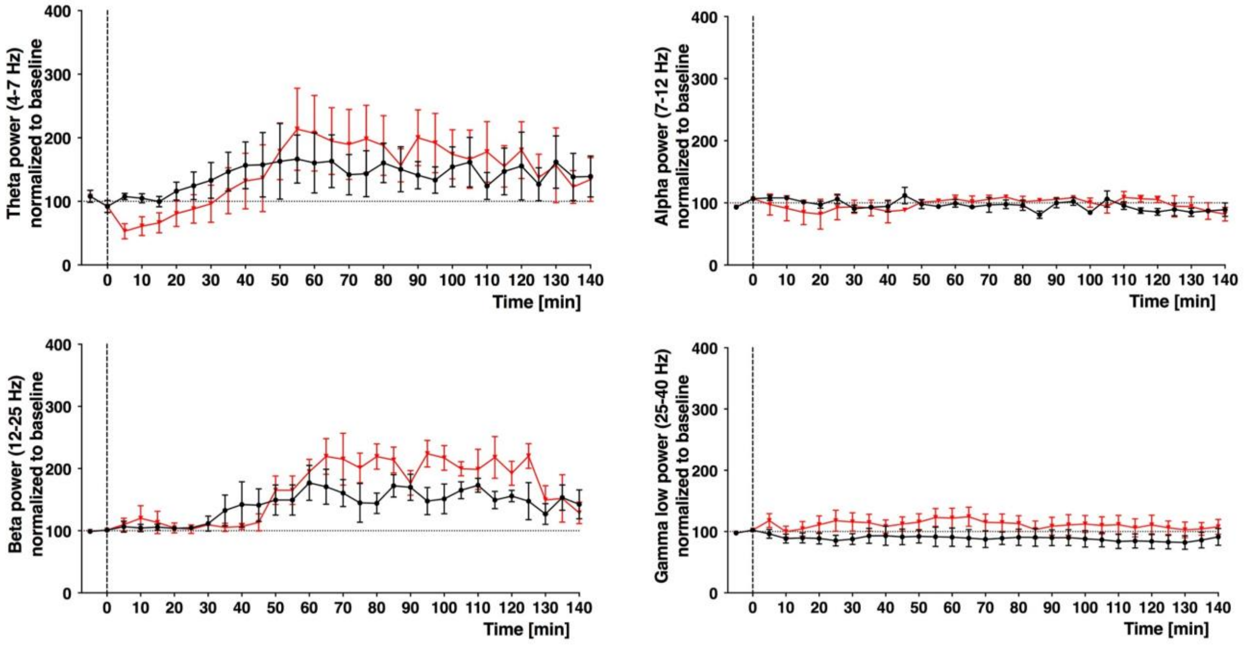


**Supplementary Figure 1.** Theta, alpha, beta and low gamma oscillations after an acute injection of saline (black) or subanesthetic ketamine (red; 10 mg/kg, i.p.). Data analyzed in 5 min bins. N=4/group.

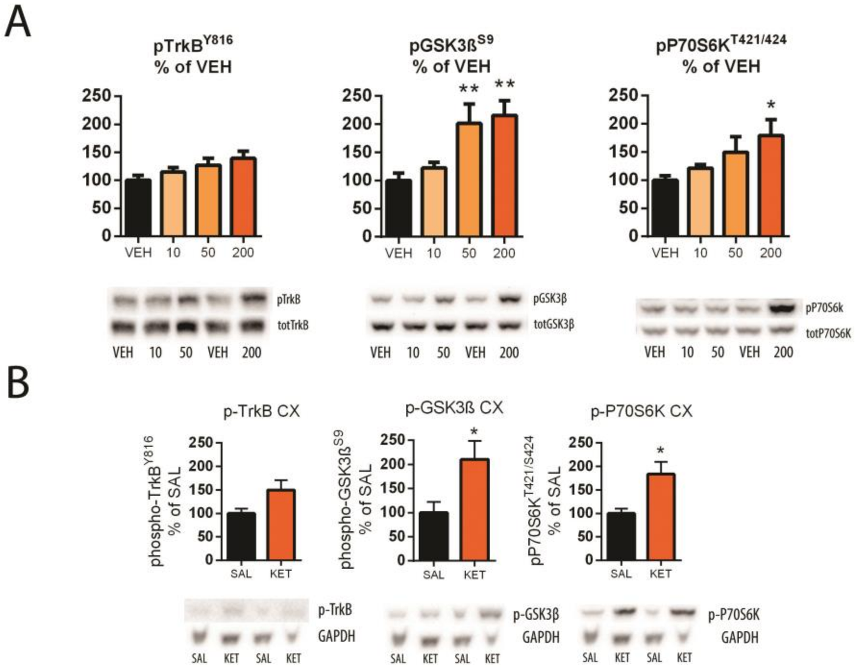


**Supplementary Figure 2.** (**A**) Phosphorylation of TrkB$^{Y816}$, GSK3β$^{S9}$ and p70S6k$^{T421/424}$ in the adult mouse medial prefrontal cortex 30-minutes after an acute i.p. injection of ketamine (10 mg/kg, 50 mg/kg, 200 mg/kg; i.p.). N=4-6/group. (**B**) Phosphorylation of TrkB$^{Y816}$, GSK3β$^{S9}$ and p70S6k$^{T421/424}$ in the adult mouse medial prefrontal cortex 3-minutes after an acute i.p. injection of high dose of ketamine (200 mg/kg; i.p.). N=5-6/group *<0.05, **<0.01, ***<0.005, one-way ANOVA followed with Dunnett´s *post hoc* test (A) or Student t-test (B).

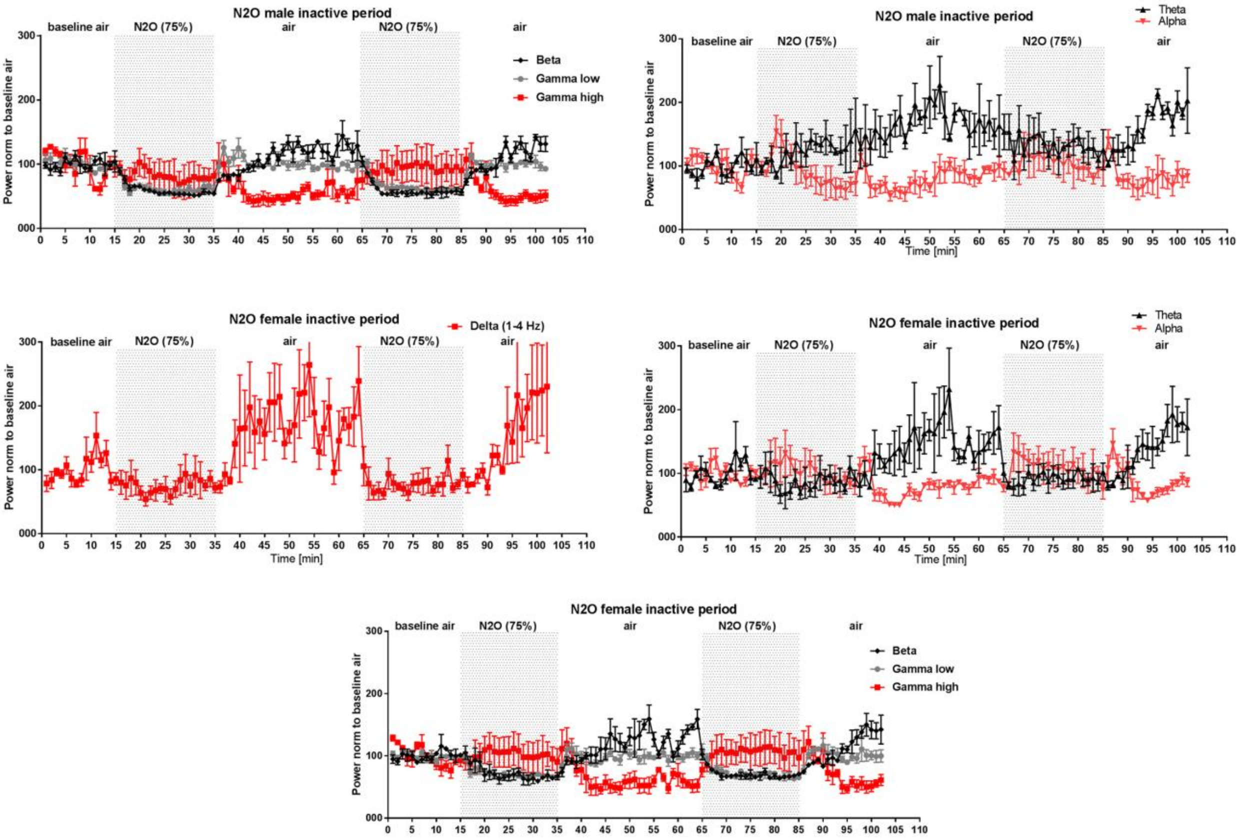


**Supplementary Figure 3.** (**A**) Normalized power of beta, gamma, theta and alpha oscillations in male mice before, during (grey background) and after $N_2O$ (75%). (**B**) Normalized power of major EEG oscillations in female mice before, during (grey background) and after $N_2O$ treatment (75%). Data analyzed in 1 min bins. N=3/group. Data are means ± S.E.M.

| *Gene* | *Forward primer* | *Reverse primer* | *Reference* |
|---|---|---|---|
| *Arc* | AAGTGCCGAGCTGAGATGC | CGACCTGTGCAACCCTTTC | Primer bank ID 9055166a1 |
| *ß-actin* | GGCTGTATTCCCCTCCATCG | CCAGTTGGTAACAATGCCATGT | Primer bank ID 6671509a1 |
| *Bdnf (exon IV)* | ACCGAAGTATGAAATAACCATAGTAAG | TGTTTACTTTGACAAGTAGTGACTGAA | Ref. 1 |
| *Bdnf (total)* | GAAGGCTGCAGGGGCATAGACAAA | TACACAGGAAGTGTCTATCCTTATG | Ref. 1 |
| *cFos* | CGGGTTTCAACGCCGACTA | TTGGCACTAGAGACGGACAGA | Primer bank ID 6753894a1 |
| *Egr1* | GCCAAGGCCGTAGACAAAATC | CCACTCCGTTCATCTGGTCA | Primer bank ID 14318592a1 |
| *Gapdh* | GGTGAAGGTCGGTGTGAACGG | CATGTAGTTGAGGTCAATGAAGGG | Ref. 2 |
| *Homer1a* | GGCAAACACTGTTTATGGACTGG | GTAATTCAGTCAACTTGAGCAACC | Ref. 3 |
| *Mkp1* | CTGCTTTGATCAACGTCTCG | AAGCTGAAGTTGGGGGAGAT | Ref. 4 |
| *Synapsin* | ACACCGACTGGGCAAAATA | GTCACAGAAGTTGTAGACAGAATG | |
| *Zif268 (egr-1)* | TCCTCTCCATCACATGCCTG | CACTCTGACACATGCTCCAG | Ref. 5 |

**Supplementary Table 1.** Primers used for quantitative RT-PCR.

| | | | | | | | |
|---|---|---|---|---|---|---|---|
| **Figure 1** | | | **n** | **Statistical test** | | **Significance** | |
| **a** | **Sham vs. flurothyl** | | n=3 | | | | |
| | EEG (delta, theta, alpha, beta, gamma) | | | No statistical testing | | | |
| **b** | **Saline vs ketamine 10 mg/kg** | | n=4 | | | | |
| | EEG (delta, high gamma) | | | No statistical testing | | | |
| **c** | **$N_2O$ 50% 1h + 1h recovery** | | n=4 | | | | |
| | EEG (delta, theta, alpha, beta, gamma) | | | No statistical testing | | | |
| **Figure 2** | | | **n** | **Statistical test** | | **Significance** | |
| **a** | **Sham vs 50% $N_2O$ 1h (1h washout)** | | | | | | |
| | *Arc* | | n = 8, 6 | Student's t-test | * | 0,0123 | |
| | *cFos* | | n = 8, 6 | Welch's t-test | * | 0,0177 | |
| | *Bdnf IV* | | n = 4, 5 | Welch's t-test | ** | 0,0083 | |
| | *Homer1* | | n = 7, 6 | Student's t-test | *** | <0,0001 | |
| | *Bdnf total* | | n = 7, 5 | Student's t-test | *** | 0,0003 | |
| | *Syn I* | | n = 7, 4 | Student's t-test | * | 0,0131 | |
| | *Zif* | | n = 8, 5 | Student's t-test | * | 0,0101 | |
| | *MKP1* | | n = 6, 4 | Student's t-test | ** | 0,0033 | |
| | *EGR2* | | n = 6, 4 | Student's t-test | *** | <0,0001 | |
| | pTrkB | | n= 6 | Student's t-test | ns | 0,4262 | |
| | pGSK3β | | n= 6 | Student's t-test | ns | 0,7957 | |
| | pP70S6K | | n= 6 | Student's t-test | ns | 0,6473 | |
| **b** | **Sham vs 50% $N_2O$ 2h (no washout) vs 50% $N_2O$ 1h (1h washout)** | | | | | | |
| | *Arc* | | n = 6,6,4 | Kruskal-Wallis one-way ANO | ** | H=10,19; p=0,0011 | |
| | | Ctrl vs. $N_2O$ 2h | | Dunn's post hoc test | * | 0,0198 | |
| | | Ctrl vs. $N_2O$ 1h/1h | | Dunn's post hoc test | ** | 0,0091 | |
| | *Bdnf IV* | | n = 6,6,4 | One-way ANOVA | * | F (2, 14) = 4,578 | P=0,0295 |
| | | Ctrl vs. $N_2O$ 2h | | Dunnett's post hoc te | * | 0,0368 | |
| | | Ctrl vs. $N_2O$ 1h/1h | | Dunnett's post hoc te | * | 0,0396 | |
| | *cFos* | | n = 6,6,4 | Kruskal-Wallis one-way ANO | *** | H=10,66; p=0,0006 | |
| | | Ctrl vs. $N_2O$ 2h | | Dunn's post hoc test | * | 0,0106 | |
| | | Ctrl vs. $N_2O$ 1h/1h | | Dunn's post hoc test | * | 0,0114 | |
| **c** | **Sham vs 50% $N_2O$ 30 min (no washout)** | | n= 6 | | | | |
| | pTrkB | | | Student's t-test | ns | 0,8668 | |
| | pGSK3β | | | Student's t-test | ns | 0,6064 | |
| | pP70S6K | | | Student's t-test | ns | 0,227 | |
| | pMAPK | | | Student's t-test | * | 0,0468 | |
| | cFos | | | Student's t-test | ** | 0,0047 | |
| **Figure 3** | | | **n** | **Statistical test** | | **Significance** | |
| **a** | **SAL vs HNK 20 mg/kg, PFC** | | n = 6/group | | | | |
| | pTrkB | | | Student's t-test | ns | 0,6967 | |
| | pGSK3β | | | Student's t-test | ns | 0,7411 | |
| | pP70S6K | | | Student's t-test | ns | 0,2945 | |
| | **SAL vs KET 10 mg/kg vs KET 100 mg/kg, PFC** | | n = 6/group | | | | |
| | pTrkB | | | Kruskal-Wallis one-way ANO | ** | H=8.573; p=0,0074 | |
| | | SAL vs KET 10mg/kg | | Dunn's post hoc test | ns | >0,9999 | |
| | | SAL vs KET 100mg/kg | | Dunn's post hoc test | * | 0,0137 | |
| | pGSK3β | | | Kruskal-Wallis one-way ANO | * | H=7,38;p=0,0181 | |
| | | SAL vs KET 10mg/kg | | Dunn's post hoc test | ns | >0,9999 | |
| | | SAL vs KET 100mg/kg | | Dunn's post hoc test | * | 0,0401 | |
| | pP70S6K | | | Kruskal-Wallis one-way ANOVA | | H=8,573; p=0,0074 | |
| | | SAL vs KET 10mg/kg | | Dunn's post hoc test | ns | >0,9999 | |
| | | SAL vs KET 100mg/kg | | Dunn's post hoc test | * | 0,0137 | |
| **b** | **SAL vs KET 100mg/kg vs d-KET 100 mg/kg, PFC** | | n = 7/group | | | | |
| | pTrkB | | | One-way ANOVA | * | F (2, 18) = 4,694 | P=0,0229 |
| | | SAL vs. KET | | Dunnett's post hoc te | ns | 0,0587 | |
| | | SAL vs. d-KET | | Dunnett's post hoc te | * | 0,018 | |
| | pGSK3β | | | One-way ANOVA | * | F (2, 18) = 4,83 | P=0,0209 |
| | | SAL vs. KET | | Dunnett's post hoc te | * | 0,0244 | |
| | | SAL vs. d-KET | | Dunnett's post hoc te | * | 0,0312 | |
| | pP70S6K | | | One-way ANOVA | * | F (2, 18) = 5,611 | P=0,0128 |
| | | SAL vs. KET | | Dunnett's post hoc te | * | 0,0308 | |
| | | SAL vs. d-KET | | Dunnett's post hoc te | * | 0,0114 | |
| **c** | **Representative time-frequency EEG spectograms** | | | no statistical testing | | | |
| **d** | **SAL vs KET 10 mg/kg vs KET 100 mg/kg vs HNK mg** | | n = 3/group | | | | |
| | Delta | | | One-way ANOVA | *** | F (3, 8) = 149 | P<0,0001 |
| | | SAL vs. K10 | | Dunnett's post hoc te | ns | 0,5881 | |
| | | SAL vs. K100 | | Dunnett's post hoc te | **** | 0,0001 | |
| | | SAL vs. HNK | | Dunnett's post hoc te | ns | 0,3819 | |
| | Theta | | | One-way ANOVA | ** | F (3, 8) = 8,225 | P=0,0079 |
| | | SAL vs. K10 | | Dunnett's post hoc te | ns | 0,8017 | |
| | | SAL vs. K100 | | Dunnett's post hoc te | * | 0,0173 | |
| | | SAL vs. HNK | | Dunnett's post hoc te | ns | 0,9406 | |
| | Alpha | | | One-way ANOVA | ns | F (3, 8) = 2,4 | P=0,1433 |
| | Beta | | | One-way ANOVA | ** | F (3, 8) = 13,84 | P=0,0016 |
| | | SAL vs. K10 | | Dunnett's post hoc te | ns | 0,5591 | |
| | | SAL vs. K100 | | Dunnett's post hoc te | ** | 0,0049 | |
| | | SAL vs. HNK | | Dunnett's post hoc te | ns | 0,7899 | |
| | Gamma low | | | One-way ANOVA | * | F (3, 8) = 6,66 | P=0,0144 |
| | | SAL vs. K10 | | Dunnett's post hoc te | ns | 0,9999 | |
| | | SAL vs. K100 | | Dunnett's post hoc te | * | 0,0162 | |
| | | SAL vs. HNK | | Dunnett's post hoc te | ns | 0,9999 | |
| | Gamma high | | | One-way ANOVA | ** | F (3, 8) = 15,39 | P=0,0011 |
| | | SAL vs. K10 | | Dunnett's post hoc te | *** | 0,0009 | |
| | | SAL vs. K100 | | Dunnett's post hoc te | ** | 0,0025 | |
| | | SAL vs. HNK | | Dunnett's post hoc te | ns | 0,2303 | |
| **Figure 4** | | | **n** | **Statistical test** | | **Significance** | |
| **a** | **Flurothyl (10 min post-seizure)** | | n=3 | | | | |
| | pTrkB | | | Student's t-test | * | 0.0226 | |
| | pGSK3β | | | Student's t-test | * | 0.0116 | |
| | pP70S6K | | | Student's t-test | *** | 0.0003 | |
| **b** | **Representative EEG spectogram** | | | no statistical testing | | | |
| **c** | **Nitrous oxide dose response (5 min washout)** | | n = 7,7,6 | | | | |
| | pTrkB | SHAM vs 50% $N_2O$ | | Student's t-test | ns | 0,3591 | |
| | | SHAM vs 65% $N_2O$ | | Student's t-test | * | 0,0112 | |
| | | SHAM vs 75% $N_2O$ | | Student's t-test | ns | 0,4079 | |
| | pGSK3β | SHAM vs 50% $N_2O$ | | Student's t-test | * | 0,041 | |
| | | SHAM vs 65% $N_2O$ | | Student's t-test | ** | 0,0023 | |
| | | SHAM vs 75% $N_2O$ | | Student's t-test | *** | 0,0001 | |
| | pP70S6K | SHAM vs 50% $N_2O$ | | Welch's t-test | ns | 0,0646 | F test * |
| | | SHAM vs 65% $N_2O$ | | Student's t-test | *** | <0,0001 | |
| | | SHAM vs 75% $N_2O$ | | Student's t-test | *** | 0,0008 | |
| **d** | **Nitrous oxide 55% (15 min washout)** | | n = 6,7 | | | | |
| | pTrkB | | | Mann-Whitney test | * | 0,014 | |
| | pGSK3β | | | Mann-Whitney test | ** | 0,0082 | |
| | pP70S6K | | | Mann-Whitney test | ** | 0,0047 | |

**Supplementary Table 2.** Statistical tests and n-numbers for main figures.
